# A Rosella-PLIN2 Knock-in Mouse Reveals Lipophagy and Immunometabolic Interplay in Atherosclerosis

**DOI:** 10.1101/2025.04.15.647781

**Authors:** Thomas Laval, Nathan Joyce, Dominique Boucher, Valérie Rochon, Christina Emerton, Mishita Dharia, Sabrina Robichaud, Victoria Lorant, My-Anh Nguyen, Michèle Geoffrion, Katey J. Rayner, Derrick Gibbings, Lauryl M. J. Nutter, Ryan C. Russell, Mireille Ouimet

## Abstract

Cytosolic lipid droplets (LDs) regulate lipid homeostasis, with abnormal LD dynamics linked to metabolic diseases like atherosclerosis. In macrophage foam cells, LDs undergo autophagic degradation via lipophagy, but the extent of this process in vascular smooth muscle cell (VSMC) foam cells remains unclear. To track lipophagy in real time, we developed a Rosella-PLIN2 biosensor by tagging perilipin 2 (PLIN2) with the fluorescent pH-biosensor Rosella. We show that proatherogenic lipoproteins and autophagy activators stimulate lipophagy in human macrophages. Targeting LDs with an LC3 fusion protein or LD-autophagy tethering compounds (LD-ATTECs) selectively enhanced lipophagy, promoting foam cell LD clearance. In an atherosclerosis model, Rosella-PLIN2 accurately tracked lipophagy in arterial foam cells, revealing distinct PLIN2 expression patterns in macrophage and non-leukocyte foam cells. We identified a lipophagy deficiency in VSMC foam cells and demonstrate that enhancing lipophagy promotes LD catabolism in primary VSMC foam cells. TREM2^+^ macrophages exhibited high lipid content and low lipophagy flux, whereas TREM2^-^ macrophages had low lipid content and high lipophagy flux. Our findings highlight a cell-specific interplay between lipophagy and immunometabolism in arterial foam cells, unveiling novel therapeutic avenues for atherosclerosis. Additionally, the Rosella-PLIN2 model provides a powerful tool for studying LD metabolism, offering new insights into lipid homeostasis and disease mechanisms.

## Introduction

Lipid droplets (LDs) are dynamic and ubiquitous organelles that store triglycerides and cholesterol esters, playing a central role in lipid metabolism. In response to environmental stressors, such as excess lipids or inflammatory signals, LDs accumulate in foam cells. Foam cells, immune-active lipid-laden cells that can profoundly reshape metabolic and inflammatory pathways, are the hallmark of atherosclerotic plaques that develop as a result of chronic inflammation and the accumulation of modified forms of cholesterol-rich low-density lipoproteins (LDL)^1,2^. Traditionally, these foam cells were thought to originate primarily from monocyte-derived macrophages^2^. However, compelling evidence from human and mouse studies challenges this dogma, revealing that vascular smooth muscle cells (VSMCs) account for 30–70% of foam cells in atherosclerotic plaques^3–7^. Intriguingly, despite their shared foam cell identity, these two cell types exhibit distinct metabolic fates for modified LDL-derived cholesterol and LDs^8^, underscoring the need to unravel their respective roles in atherosclerosis.

A promising strategy to mitigate cholesterol accumulation in plaques is to enhance cholesterol efflux from foam cells, the rate-limiting step of reverse cholesterol transport (RCT) - a potent anti-atherogenic process that facilitates cholesterol excretion via high-density lipoproteins (HDL)^9^. This process hinges on the release of free cholesterol from LDs, mediated by neutral cytoplasmic lipases or lysosomal acid lipase (LAL) following LD delivery to lysosomes^10^. Autophagy, a degradative pathway that traffics intracellular cargo to lysosomes, plays a crucial role in regulating macrophage LD breakdown and cholesterol efflux^10^. In macrophage foam cells, autophagy-mediated cholesterol efflux occurs through a selective process known as lipophagy, where LDs are tethered to autophagosomes via microtubule-associated protein 1 light chain 3 (LC3) and delivered to lysosomes^11,12^. Notably, several candidate lipophagy factors have been identified, many of which become dysregulated during atherogenesis, linking impaired lipophagy to disease progression^11,13–15^.

Autophagy flux declines as atherosclerosis advances^6,16^, with an even greater impairment observed in VSMC-derived foam cells compared to their macrophage counterparts^6^. However, critical questions remain, such as whether all plaque foam cells form *bona fide* LDs and how lipophagy is selectively regulated across distinct foam cell populations^17^. To this end, we developed a Rosella-PLIN2 lipophagy reporter by tagging the ubiquitous LD coat protein perilipin 2 (PLIN2)^18^ with the pH-sensitive biosensor Rosella^19^. Leveraging this novel lipophagy reporter and selective autophagy enhancers, we show that lipophagy regulates macrophage foam cell immunometabolism. Furthermore, we introduce the Rosella-PLIN2 knock-in mouse – the first whole-body lipophagy reporter model – offering unprecedented insights into LD metabolism *in vivo*.

In a model of atherosclerosis, we show that macrophage and non-leukocyte foam cells exhibit distinct patterns of LD accumulation, accompanied by differential PLIN2 expression. Furthermore, we provide strong evidence that enhancing lipophagy drives LD catabolism in primary VSMC foam cells. We also identify foam cell subpopulations within atherosclerotic lesions that exhibit varying levels of lipophagy activity, such as Lipid^hi^Lipophagy^lo^TREM2^+^ macrophages. Our findings uncover a cell-specific interplay between lipophagy and immunometabolism in arterial foam cells, while the Rosella-PLIN2 model provides a powerful tool for advancing the study of LD metabolism in health and disease.

## Results

### A new lipophagy biosensor

Several biosensors have been developed to monitor autophagy in live cells non-invasively, including Rosella, which leverages pH changes during autophagy^19^. Rosella consists of a fusion between the pH-stable DsRed.T2 pH-sensitive pHluorin. It fluoresces both red and green in neutral conditions, while in acidic lysosomes, green fluorescence is quenched, leaving only red fluorescence^19^. This enables tracking of autophagic cargo degradation via fluorescence microscopy and flow cytometry. Perilipins (PLINs) are key LD structural proteins, with PLIN2 being ubiquitously expressed^18^ and the predominant LD coat protein in macrophage foam cells^10^. To selectively track lipophagy, we generated a Rosella-PLIN2 fusion protein reporter by tagging PLIN2 at its N-terminus with Rosella (**Fig.1a**). Transient expression in oleic acid-treated HEK293 cells confirmed that Rosella-PLIN2 localized to cytosolic LDs, while rapamycin-induced autophagy increased lysosomal red fluorescence, validating its biosensor function (**Fig.1b**).

**Fig.1:**
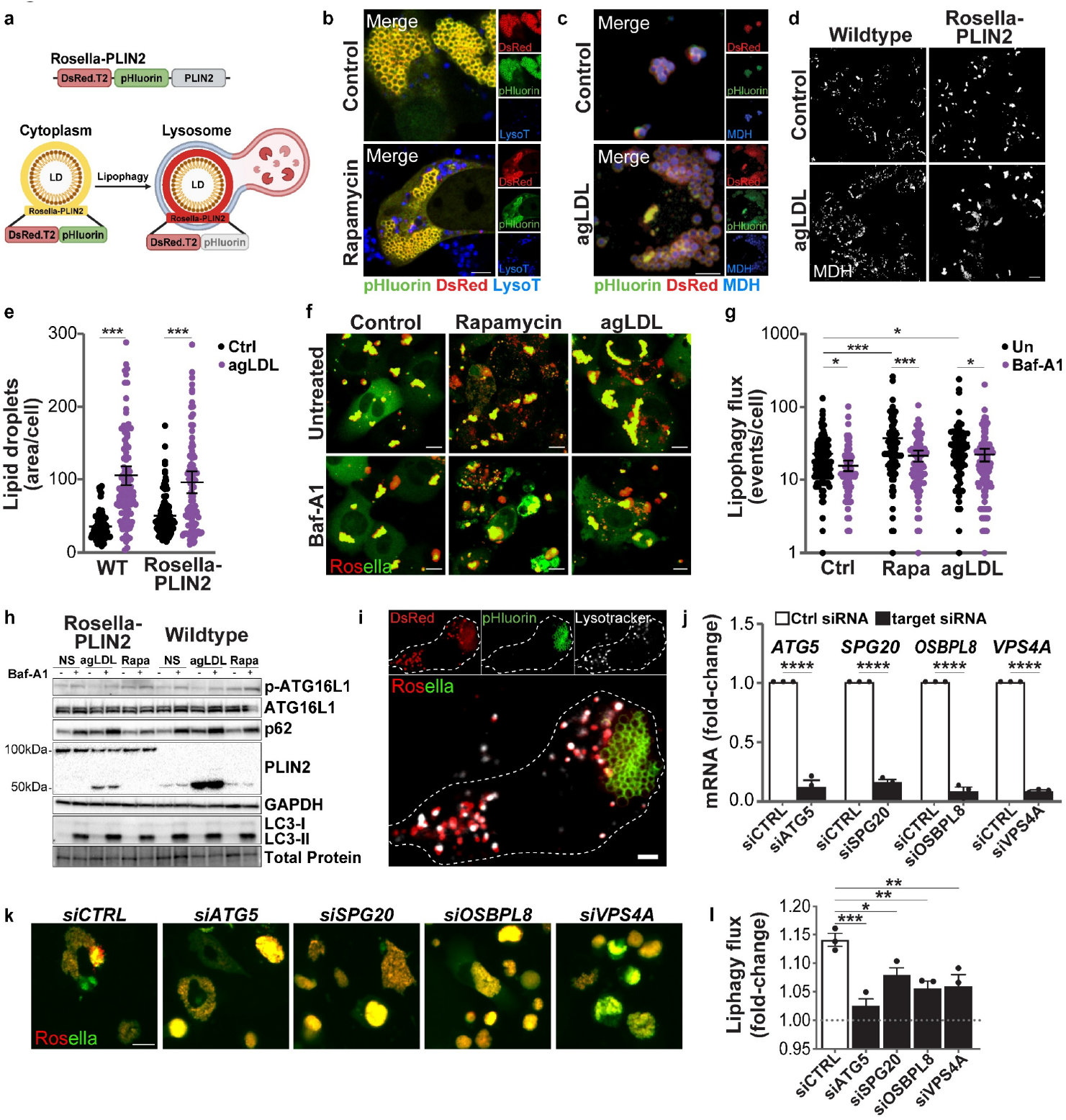
The Rosella-PLIN2 lipophagy reporter. (**a**) Schematic of the Rosella-PLIN2 lipophagy biosensor. (**b**) Live-cell imaging of HEK293T cells expressing Rosella-PLIN2, loaded with oleic acid for 24h, and treated or not with rapamycin for 3h, then stained with Lysotracker (LysoT). Sale bar: 5μm. (**c**) Fluorescence microscopy of Rosella-PLIN2 THP-1 macrophages, incubated or not with aggregated LDL (agLDL) for 24h, then stained with monodansylpentane (MDH). Scale bar: 5μm. (**d-e**) Representative images and quantification of neutral lipid content in wildtype and Rosella-PLIN2 THP-1 macrophages. (n>88 cells per condition). Scale bar: 20μm. (**f-g**) Live-cell imaging and quantification of lipophagy in Rosella-PLIN2 THP-1 macrophages treated with rapamycin or agLDL for 24h, with or without bafilomycin A1 (Baf-A1) for the last 2h. (n>75 cells per condition). Scale bar: 10μm. (**h**) Immunoblot analysis of wildtype and Rosella-PLIN2 THP-1 macrophages treated as in (**f**). (**i**) Live-cell imaging of oleic acid-loaded Rosella-PLIN2 THP-1 macrophages treated with Torin-1 for 24h and stained with LysoTracker. Scale bar: 2μm. (**j**) mRNA expression levels of target genes in Rosella-PIN2 THP-1 macrophages 48h post-transfection with non-targeting (siCTRL) or targeting siRNAs. (n=3 independent experiments) (**k-l**) Live-cell imaging and quantification of lipophagy in siRNA-transfected Rosella-PLIN2 THP-1 cells loaded with oleic acid and treated with Torin-1 or vehicle for 24h, with representative images of Torin-1-treated cells. Lipophagy flux is expressed as a fold change of Torin-1 relative to vehicle. (n=3 independent experiments, >150 cells per condition). Scale bar: 20µm. Data are mean ± 95%CI, two-way ANOVA with Holm-Sidak post-hoc multiple comparison tests (**e, g**), or mean ± SEM, Student’s *t*-test (**j**) or one-way ANOVA with Dunnett post-hoc multiple comparison test (**l**).

Next, we established a stably expressing Rosella-PLIN2 THP-1 monocytic cell line. As expected, Rosella-PLIN2 localized to the surface of LDs in differentiated THP-1 macrophages (**Fig.1c**). Upon aggregated LDL (agLDL) treatment, Rosella-PLIN2 THP-1 macrophages accumulated neutral lipids comparably to wildtype cells (**Fig.1d-e**). Rapamycin or agLDL treatment increased red-only puncta in lysosomes, which were restored to dual fluorescence upon bafilomycin A1 (Baf-A1) treatment, confirming acidification-dependent fluorescence quenching of the Rosella-PLIN2 biosensor (**Fig.1f**). Quantification of lipophagy flux showed a significant increase with rapamycin and, to a lesser extent, agLDL, while Baf-A1 treatment suppressed it (**Fig.1g**). Rosella-PLIN2 was expressed at the expected molecular weight without interfering with endogenous PLIN2 upregulation upon agLDL loading (**Fig.1h**). Autophagy flux, assessed by LC3-I/LC3-II, p62/SQSTM1 and phospho-ATG16L1 (pATG16L1) levels, remained comparable between wildtype and Rosella-PLIN2 THP-1 macrophages (**Fig.1h**).

Like rapamycin, the mTOR inhibitor Torin-1 increased lysosome-associated lipophagy events in oleic acid-loaded Rosella-PLIN2 THP-1 macrophages (**Fig.1i**). Recent studies implicate SPG20^15^, ORP8^13^ and VPS4A^14^ in selective LD turnover. siRNA-mediated knockdown of ATG5, SPG20, ORP8 and VPS4A in Rosella-PLIN2 THP-1 macrophages reduced target mRNA by >80% (**Fig.1j**). Torin-1 treatment increased lipophagy in oleic acid-loaded Rosella-PLIN2 THP-1 macrophages treated with control siRNA, compared to the vehicle-alone control (**Fig.1k-l**). In contrast, ATG5 knockdown impaired Torin-1-induced lipophagy, confirming its reliance on core autophagy machinery (**Fig.1k-l**). Similarly, knockdown of SPG20, VPS4A, and OSBPL8 reduced lipophagy activation (**Fig.1k-l**), supporting their role in macrophage lipophagy regulation. Collectively, these results validate the functionality of our generated lipophagy biosensor.

### Selective activation of lipophagy promotes macrophage LD catabolism

A recent advancement in the field of lipophagy research was the development of the first specific lipophagy-inducing compounds, achieved by linking LD- and LC3-interacting molecules (**Fig.2a**). These LD-autophagy tethering compounds (LD-ATTECs) recruit LC3 to LDs, thereby promoting lipophagic LD degradation^20^. To assess the ability of LD-ATTECs to reduce LD accumulation in macrophage foam cells, we treated agLDL-loaded Rosella-PLIN2 THP-1 macrophage foam cells with LD-ATTECs. This treatment significantly increased lipophagy events compared to controls (**Fig.2b-c**). Increased lipophagy flux in agLDL-loaded bone marrow-derived macrophages (BMDMs) by LD-ATTECs resulted in a marked decrease in cytosolic LDs (**Fig.2d-e**), which could be blocked by the lysosome acid lipase (LAL) inhibitor lalistat (**Fig.2f**), confirming that lipophagy promotes LD degradation.

**Fig.2:**
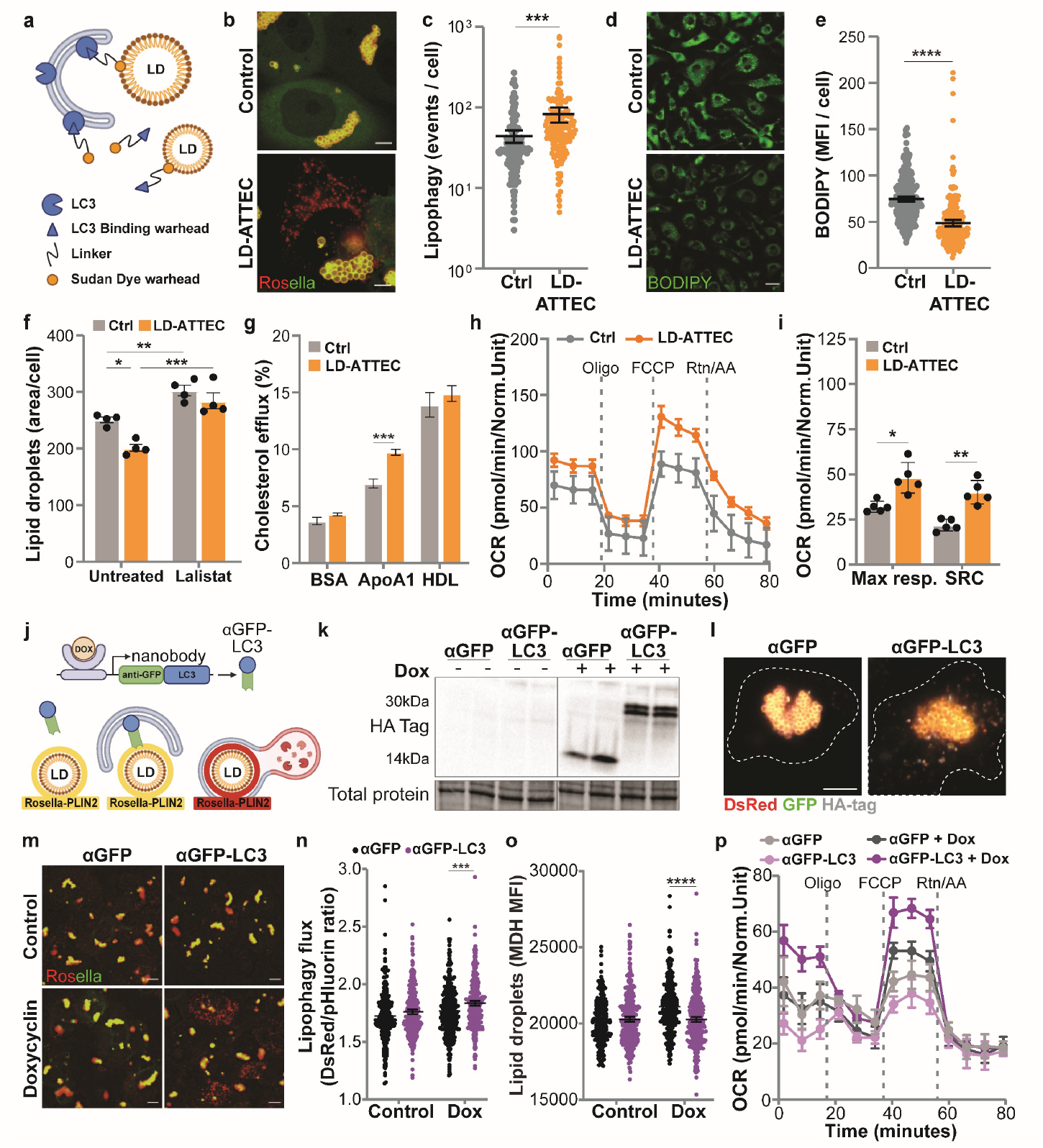
Enhancing selective lipophagy stimulates lipid catabolism and cholesterol efflux in macrophage foam cells. (**a**) Schematic of LD-ATTECs. (**b-c**) Representative images and quantification of lipophagy in agLDL-loaded Rosella-PLIN2 THP-1 macrophages treated with LD-ATTECs or vehicle (Ctrl) for 24h. (n>192 cells per condition). Scale bar: 5µm. (**d-e**) Representative images and quantification of BODIPY fluorescence in agLDL-loaded BMDMs treated as in (**b**) for 24h. (n>132 cells per condition). Scale bar: 20µm. (**f**) Lipid droplet content in agLDL-loaded BMDMs treated with LD-ATTECs ± Lalistat-2 for 24 h. (n=3 technical replicates, >259 cells per condition). (**g**) Cholesterol efflux to ApoA1 and HDL in BMDMs treated as in (**d**) for 24h. (n=3 biological replicates). (**h-i**) Oxygen Consumption Rate (OCR) (**h**) and Spare Respiratory Capacity (SRC) (**i**) in wildtype THP-1 treated as in (**b**) for 24 h. (n=5 technical replicates). (**j**) Schematic of anti-GFP nanobody (αGFP)-LC3 chimera. (**k**) Immunoblot analysis of αGFP-Ctrl and αGFP-LC3 THP-1 macrophages treated ± doxycycline (Dox) for 24h and incubated with Baf-A1 for 2h. (**l**) Nanobody localization to lipid droplets in αGFP-Ctrl and αGFP-LC3 THP-1 macrophages treated with doxycycline for 24h. Scale Bar: 5um. (**m-o**) Live-cell images and quantification of lipophagy and MDH fluorescence in αGFP-Ctrl and αGFP-LC3 THP-1 macrophages loaded with OA and treated ± doxycycline for 24h. (n>222 cells per condition). Scale bar: 10µm. (**p**) OCR in αGFP-Ctrl and αGFP-LC3 THP-1 macrophages treated as in (**m**). (n=5 technical replicates). Data are mean ± 95%CI (**c, e, n, o**), mean ± SEM (**f-i, p**), Student *t*-test (**c, e, i**) or two-way ANOVA with Tukey (**f**) or Sidak (**g, n, o**) post-hoc multiple comparison tests.

Within lysosomes, the hydrolysis of LD cholesterol esters by LAL generates free cholesterol, which can be exported via ABCA1-mediated efflux^10^. In parallel, free fatty acids derived from LD triglycerides and/or cholesterol esters through lipophagy can be channeled to mitochondria for β-oxidation^12^. In line with this, LD-ATTECs enhanced cholesterol efflux to lipid-poor apoA1 (**Fig.2g**) and increased oxygen consumption, indicating elevated mitochondrial respiration in LD-ATTEC-treated cells (**Fig.2h-i**).

To further investigate the potential of selective lipophagy activation to reverse lipid accumulation in macrophage foam cells, we utilized anti-GFP nanobodies fused to LC3 (αGFP-LC3) to target Rosella-PLIN2 LDs and promote autophagosome biogenesis at their surface (**Fig.2j**). We generated Rosella-PLIN2-expressing THP-1 cell lines stably expressing either αGFP alone or αGFP-LC3 under the control of a doxycycline-inducible promoter. As expected, doxycycline treatment significantly increased the expression of both αGFP and αGFP-LC3 nanobodies (**Fig.2k**). Both the αGFP and αGFP-LC3 are predicted to recognize the pHluorin (GFP) domain of Rosella-PLIN2^21^, but only αGFP-LC3 should initiate autophagosome biogenesis at the LD surface. We confirmed that αGFP and αGFP-LC3 localized to Rosella-PLIN2-tagged LDs, indicating successful targeting of the Rosella-PLIN2 pHluorin domain (**Fig.2l**). Notably, αGFP-LC3 expression upon doxycycline treatment led to a substantial increase in lipophagy events compared to cells expressing αGFP alone (**Fig.2m-n**). This was accompanied by a concomitant reduction in cellular LD content in cells expressing αGFP-LC3 as compared to αGFP alone (**Fig.2o**). Furthermore, oleic acid (OA)-loaded macrophages exhibited increased oxygen consumption following doxycycline-induced αGFP-LC3 expression compared to control (**Fig.2p**), reinforcing the findings observed with LD-ATTECs. Together, these results provide strong evidence that selective lipophagy activation, through either LD-ATTECs or αGFP-LC3 nanobodies, effectively promotes LD degradation, cholesterol efflux, and enhanced mitochondrial respiration in macrophage foam cells.

### Enhancing lipophagy promotes an anti-inflammatory macrophage phenotype

Toll-like receptor (TLR) signaling is implicated in macrophage LD accumulation within human atherosclerotic plaques^22^. Furthermore, TLR stimulation can regulate both macrophage autophagy and LD formation^23–26^, suggesting that lipophagy may play a role in TLR-dependent modulation of macrophage LD metabolism. To investigate this, we stimulated Rosella-PLIN2 THP-1 macrophages for 24h with TLR2 ligands Pam3Csk4 and lipoteichoic acid from *S. aureus* (LTA), as well as the TLR4 ligand lipopolysaccharide (LPS). Notably, TLR2 ligands, but not LPS, significantly increased lipophagy, reaching levels comparable to those induced by Torin-1 (**Fig.3a**). Unlike Torin-1, however, TLR2 ligands did not enhance general autophagy flux at 24h (**Fig.3b**), suggesting a selective effect on macrophage lipophagy.

**Fig.3:**
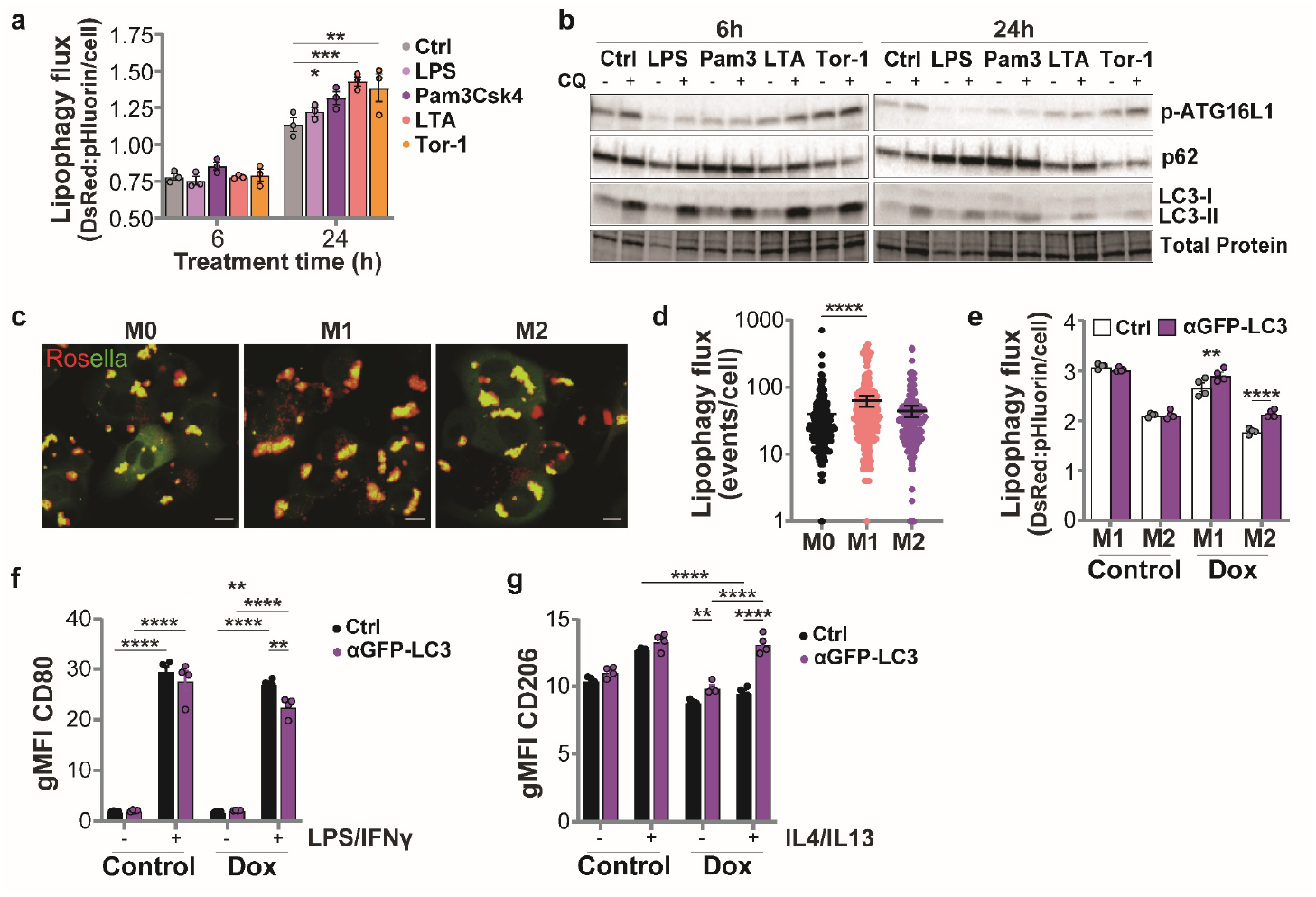
Lipophagy regulation and macrophage polarization. (**a**) Quantification of lipophagy in Rosella-PLIN2 THP-1 cells treated with lipopolysaccharide (LPS), Pam3Csk4 (Pam3), lipoteichoic acid (LTA), or Torin-1 for 6 or 24 h. (n=3 independent experiments, >150 cells per condition). (**b**) Immunoblotting of THP-1 treated as in (**a**) and treated with chloroquine (CQ) for 2h. (**c-d**) Live-cell imaging and lipophagy quantification in oleic acid-loaded Rosella-PLIN2 THP-1 cells, untreated (M0) or treated with LPS/IFNγ (M1) or IL-4/IL-13 (M2) for 24h. (n>154 cells per group). Scale bar: 10μm. (**e-g**) Lipophagy quantification (**e**) and flow cytometry analysis of CD80/CD206 expression (**f-g**) on αGFP-Ctrl and αGFP-LC3 Rosella-PLIN2 THP-1 cells, treated as in (**c**) with or without doxycycline for 24h. (n=4 technical replicates). Data are mean ± SEM, two-way ANOVA with Dunnett (**a**), Sidak (**e**) or Tukey (**f, g**) post-hoc multiple comparison test, or mean ± 95%CI, one-way ANOVA with Dunnet post-hoc multiple comparison tests (**d**).

Given the known role of LD dynamics in macrophage immune responses^27,28^, our observation that lipophagy is modulated by TLR signaling in macrophage foam cells suggests a potential crosstalk between macrophage polarization and lipophagy. Supporting this, Rosella-PLIN2 THP-1 macrophages exhibited increased lipophagy events following M1-like polarization but not M2-like polarization (**Fig.3c-d**). To further investigate this, we induced the expression of αGFP-LC3 in Rosella-PLIN2 THP-1 macrophages preloaded with oleic acid and polarized them toward M1 or M2 phenotypes, using CD80 and CD206 as respective markers (**Fig.3e-g**). Doxycycline treatment increased lipophagy flux in αGFP-LC3 but not αGFP expressing M1 and M2 Rosella-PLIN2 macrophages (**Fig.3e**). Upon doxycycline treatment, αGFP-LC3 M1-like macrophages displayed lower CD80 expression compared to αGFP-Ctrl (**Fig.3f**), while αGFP-LC3 M2-like macrophages exhibited upregulated CD206 expression, with a more modest increase observed in M0 macrophages (**Fig.3g**). These findings reveal that lipophagy stimulation promotes an anti-inflammatory macrophage phenotype.

### Generation of Rosella-PLIN2 biosensor knock-in mice to track lipophagy *in vivo*

To investigate the regulation of lipophagy during atherosclerosis development, we generated the first-ever lipophagy reporter mouse, leveraging our validated Rosella-PLIN2 biosensor. Using a knock-in strategy previously employed to generate tdTom-Plin2 mice without histological alterations^29^, we inserted the *Rosella* gene upstream of the endogenous *Plin2* locus with a short linker sequence (**Fig.4a**). This innovative design enables real-time tracking of lipophagy *in vivo*. As expected, Rosella-PLIN2 fluorescence was readily detectable in the livers of homozygous Rosella-PLIN2 mice (**Fig.4b**). Additionally, we observed enhanced Rosella-PLIN2 expression in mice fed a Western diet compared to those on a chow diet (**Fig.4b**). This was also confirmed by Western blot analysis, which revealed significantly elevated hepatic Rosella-PLIN2 levels in mice on a Western diet compared to those on chow (**Fig.4c**). Notably, the magnitude of this increase was comparable to that of endogenous PLIN2 in wildtype mice (**Fig.4c**).

**Fig.4:**
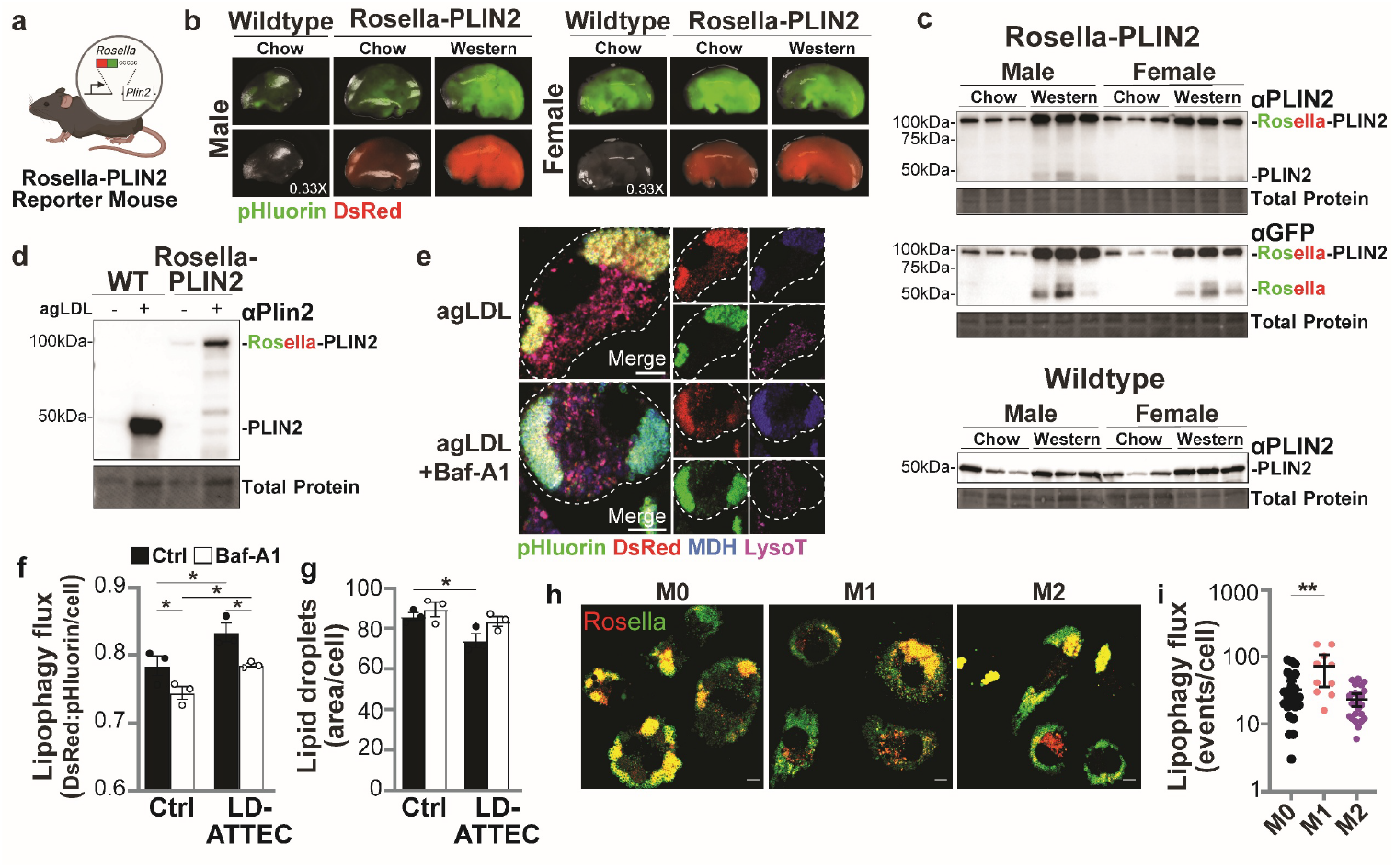
Tracking lipophagy in mice and macrophage foam cells using the Rosella lipid droplet biosensor. (**a**) Schematic of the Rosella knock-in upstream of endogenous mouse *Plin2*. (**b**) Fluorescence imaging of livers from Wildtype or Rosella-PLIN2 mice on a chow or Western diet for 5 weeks following AAV-PCSK9 injection. (**c**) Immunoblot analysis of liver homogenates from Wildtype or Rosella-PLIN2 mice fed chow or Western diet for 12 weeks following AAV-PCSK9 injection. (**d**) Immunoblot analysis of agLDL-loaded BMDMs from wildtype (WT) or Rosella-PLIN2 mice. (**e**) Representative live-cell images of agLDL-loaded Rosella-PLIN2 BMDMs treated with or without Baf-A1 for 2h, followed by MDH and LysoTracker staining. Scale bar: 5μm. (**f-g**) Quantification of lipophagy and lipid droplet content in agLDL-loaded Rosella-PLIN2 BMDMs, with or without LD-ATTEC treatment (24h) and Baf-A1 (2h) in ApoA1-supplemented lipid-free media. (n=3 technical replicates, >180 cells per condition). (**h-i**) Representative images and quantification of lipophagy in agLDL-loaded Rosella-PLIN2 BMDMs incubated with LPS/IFNγ (M1), IL-4 (M2) or left untreated (M0) for 24h. (n=10 or 24 cells per group). Scale bar: 5μm. Data are mean ± SEM, two-way ANOVA with Fisher’s LSD test (**f-g**), or mean ± 95%CI, one-way ANOVA with Dunnett post-hoc multiple comparison tests (**i**).

To validate the functionality of the Rosella-PLIN2 biosensor in macrophages, we isolated BMDMs from Rosella-PLIN2 mice and exposed them to agLDL. Rosella-PLIN2 expression was significantly increased in agLDL-loaded BMDMs compared to their unloaded counterparts (**Fig.4d**). As expected, Rosella-tagged LDs accumulated in these cells, and we observed distinct red-only puncta co-localized with lysosomes (**Fig.4e**), providing direct evidence of lipophagy. The pH sensitivity of the Rosella-PLIN2 reporter was further confirmed by treating cells with Baf-A1, which partially restored pHluorin fluorescence within lysosome-localized puncta (**Fig.4e**).

To assess biosensor functionality under enhanced lipophagy conditions, we treated agLDL-loaded BMDMs with LD-ATTECs in apoA1-supplemented media. Notably, LD-ATTEC treatment significantly increased lipophagy while simultaneously reducing LD content compared to controls (**Fig.4f**), confirming that lipophagy activity is fully preserved in Rosella-PLIN2 macrophages. In line with our observations in THP-1 macrophages, M1-like polarization enhanced lipophagy in primary Rosella-PLIN2 BMDMs compared to M0 and M2 phenotypes (**Fig.4g**), reinforcing the link between macrophage activation states and lipophagy regulation. Overall, our results establish the Rosella-PLIN2 knock-in mouse as a novel tool for studying lipophagy dynamics *in vivo*. Consistent with findings in tdTom-Plin2 mice^29^, our data confirm that endogenous Rosella tagging of *Plin2* does not alter LD metabolism, making this model a powerful new resource for investigating the role of lipophagy in cardiometabolic diseases.

### Vascular smooth muscle cells and macrophages differentially accumulate PLIN2+ lipid droplets in atherosclerotic lesions

VSMCs constitute 30-70% of foam cells in human and murine atherosclerotic lesions^3–7^. As VSMCs transdifferentiate into macrophage-like foam cells, they acquire CD68 but lack the pan-leukocyte marker CD45, distinguishing them from macrophage foam cells^4,5,7,30^. To investigate whether VSMC foam cells store lipids in *bona fide* LDs *in vivo*, we used Rosella-PLIN2 knock-in mice as reporters of PLIN2 expression. Hypercholesterolemia was induced in both wildtype and Rosella-PLIN2 mice through AAV8-gain-of-function PCSK9 overexpression and Western diet feeding for 12 weeks.

Macrophage (CD68^+^CD45^+^) and VSMC (CD68^+^CD45^-^) foam cells were identified in similar proportions in the aortic sinuses of both Rosella-PLIN2 and wildtype mice, across both sexes (**Fig.5a-c**). Notably, very few CD68^-^ plaque cells expressed Rosella-PLIN2, whereas nearly half of non-leukocyte foam cells and ∼60% of macrophage foam cells were Rosella-PLIN2-positive, consistent with CD68 as a marker for foam cells in atherosclerotic lesions (**Fig5.c-d**). Both macrophages and VSMCs showed comparable levels of pHluorin and DsRed fluorescence intensities, detectable by widefield microscopy well above tissue autofluorescence levels (**Fig.5e-f**). pHluorin and DsRed MFI was comparable between foam cell subtypes among Rosella^+^ cells, indicating similar PLIN2 expression levels (**Fig.5g**).

**Fig.5:**
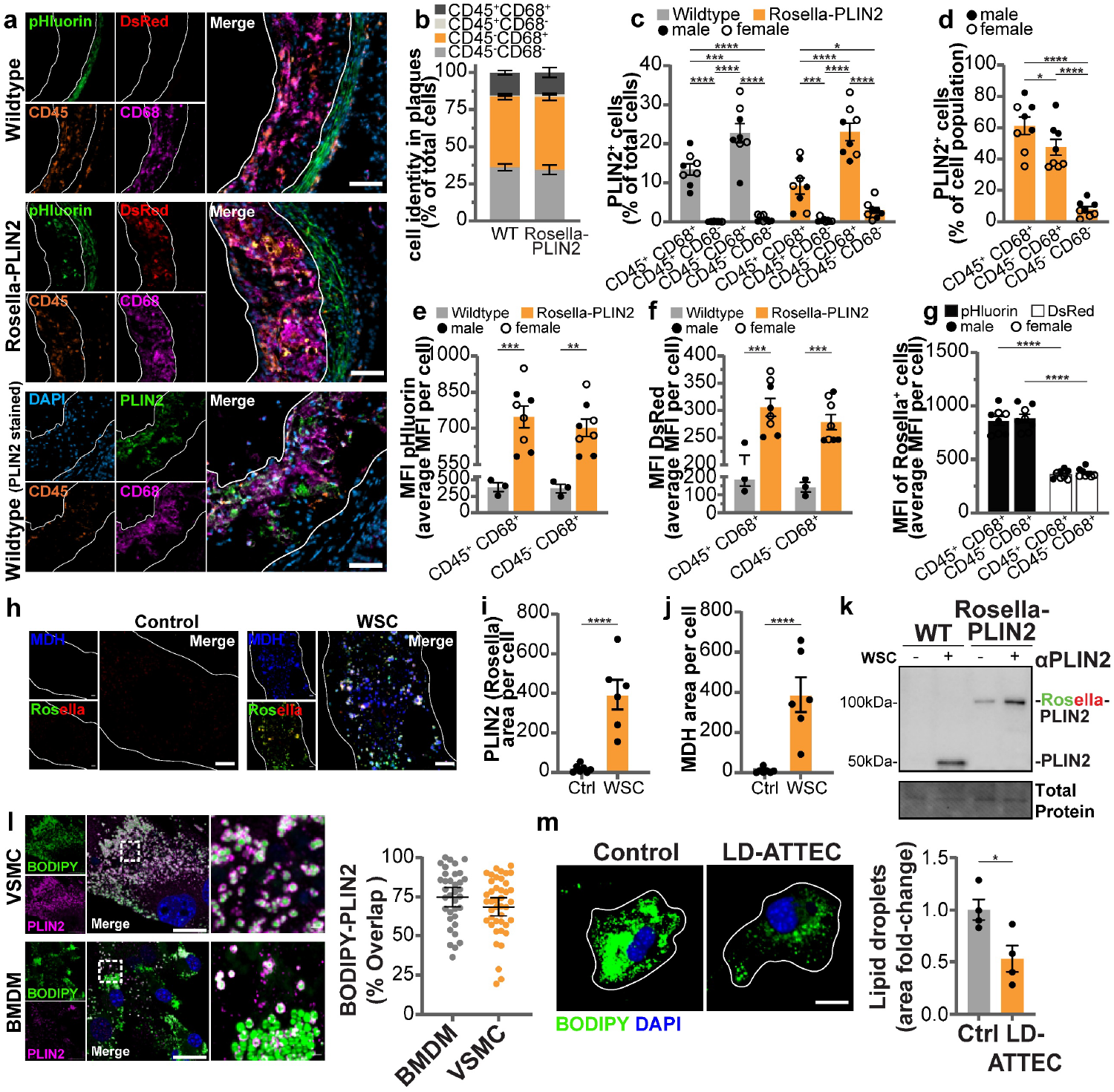
Leukocyte and non-leukocyte foam cells differentially accumulate PLIN2^+^ LDs during atherosclerosis. (**a**) Representative immunofluorescence staining of CD68 (purple), CD45 (orange), and DAPI (blue), with endogenous DsRed (red) and pHluorin (green) fluorescence in aortic sinus atherosclerotic plaques from wildtype and Rosella-PLIN2 mice. Immunostaining of CD68 (purple), CD45 (orange), DAPI (blue), and PLIN2 (green) in sinus sections from wildtype mice. Scale bar: 50μm. (**b**) Expression of CD45 and CD68 as a percentage of total plaque cells in wildtype and Rosella-PLIN2 atherosclerotic plaques. (n=3-4 mice per sex pooled). (**c**) PLIN2^+^ cells, grouped by CD45 and CD68 as a percentage of total plaque cells in wildtype (PLIN2 stained) and Rosella-PLIN2 atherosclerotic plaques. (n=3-4 mice per sex pooled, females=open circles). (**d**) PLIN2^+^ cells as a percent of respective cell populations in Rosella-PLIN2 plaques. (**e-f**) Mean fluorescence intensity (MFI) of pHluorin and DsRed in CD45^+^CD68^+^ and CD45^-^CD68^+^ cells. (**g**) MFI of pHluorin and DsRed in Rosella^+^CD45^+^CD68^+^ and Rosella^+^CD45^-^CD68^+^ cells. (**h-j**) Representative images (**h**) and quantification of Rosella-tagged PLIN2 (**i**) and MDH areas (**j**) in VSMCs from Rosella-PLIN2 mice loaded with WSC or left untreated (Ctrl) for 48 h. (n>6 cells per condition). Scale bar: 10μm. (**k**) Immunoblotting of wildtype or Rosella-PLIN2 VSMCs treated as per (**i-j**). (**l**) Representative images and quantification of BODIPY-PLIN2 area overlap in wildtype VSMCs and BMDMs loaded with WSC for 48h. (n>35 cells per condition). Scale bar: 20μm. (**m**) Representative images and quantification of BODIPY area in WSC-loaded wildtype VSMCs treated with LD-ATTEC or vehicle control (Ctrl) for 24 h in ApoA1-supplemented lipid-free media. (n=3 independent experiments, >27 cells per condition). Data are mean ± SEM or mean ± 95%CI (**l**), one- (**d**) or two-way ANOVA with Tukey (**c**) or Sidak (**d-g**) post-hoc multiple comparison tests, or Student’s *t*-test (**i-j, l-m**).

While a large proportion of VSMC foam cells formed PLIN2^+^ LDs *in vivo*, a substantial number of CD45^-^CD68^+^ foam cells lacked detectable PLIN2, suggesting that these cells store lipids either outside *bona fide* LDs or in LDs coated with other perilipins, as has been observed in other cell types^31^. Analysis of PLIN2^+^ droplets in VSMC and macrophage foam cells from wildtype mice using an anti-PLIN2 antibody yielded results consistent with the Rosella-PLIN2 mice (**Fig.5c**), confirming that Rosella-PLIN2 accurately tracks PLIN2 expression in distinct foam cell populations *in vivo*.

### Lipophagy facilitates the degradation of LDs in VSMCs

We next examined the capacity of VSMCs to degrade LDs through lipophagy. Primary VSMCs derived from Rosella-PLIN2 mice were loaded with water-soluble cholesterol (WSC) to convert them to macrophage-like foam cells and to assess their ability to accumulate LDs. WSC treatment resulted in a significant increase in neutral lipid content and a notable upregulation of Rosella-PLIN2 expression on LDs in VSMCs compared to controls (**Fig.5h-k**). PLIN2 colocalized with neutral lipids in wildtype VSMCs loaded with WSC, a pattern that was also observed in WSC-loaded BMDMs (**Fig.5l**). We then tested whether VSMC foam cells could degrade these *bona fide* LDs upon lipophagy stimulation using LD-ATTECs. LD-ATTECs significantly reduced neutral lipid content in WSC-loaded VSMCs (**Fig.5m**). Our data demonstrates that VSMC foam cells can form PLIN2^+^ LDs in the lipid-rich microenvironment of atherosclerotic plaques as well as *in vitro* following cholesterol loading and possess the capacity to degrade these LDs through lipophagy.

### Impaired lipophagy in TREM2^+^ and pro-inflammatory macrophage foam cells

To quantify lipophagy flux in foam cell populations, we analyzed Rosella-expressing cells isolated from the aortic arches of 12-week Western diet-fed Rosella-PLIN2 mice using flow cytometry. As previously^6^, aortic foam cells were identified based on their granularity and neutral lipid content, while Rosella^+^ cells were gated according to their pHluorin and DsRed fluorescence (**Fig.6a**). Confirming our gating strategy, Rosella^+^ cells represented 55.8% of foam cells and 2.7% of non-foam cells in the aortic arches of Rosella-PLIN2 mice (**Fig.6b**). Importantly, the proportions of major atheroma cell types were similar within both total foam cells and Rosella^+^ cells, with less than 50% of either population being leukocytes on average (**Fig.6c**), consistent with previous reports^4,6^. Notably, Rosella expression was observed in more than 85%, 45%, and 50% of macrophage, endothelial, and CD31^-^CD45^-^ (primarily VSMCs and fibroblasts) foam cells, respectively (**Fig.6d**). These findings, supported by our aortic sinus data in **Fig.5**, indicate that PLIN2 is preferentially expressed in macrophages within atherosclerotic lesions.

**Fig.6:**
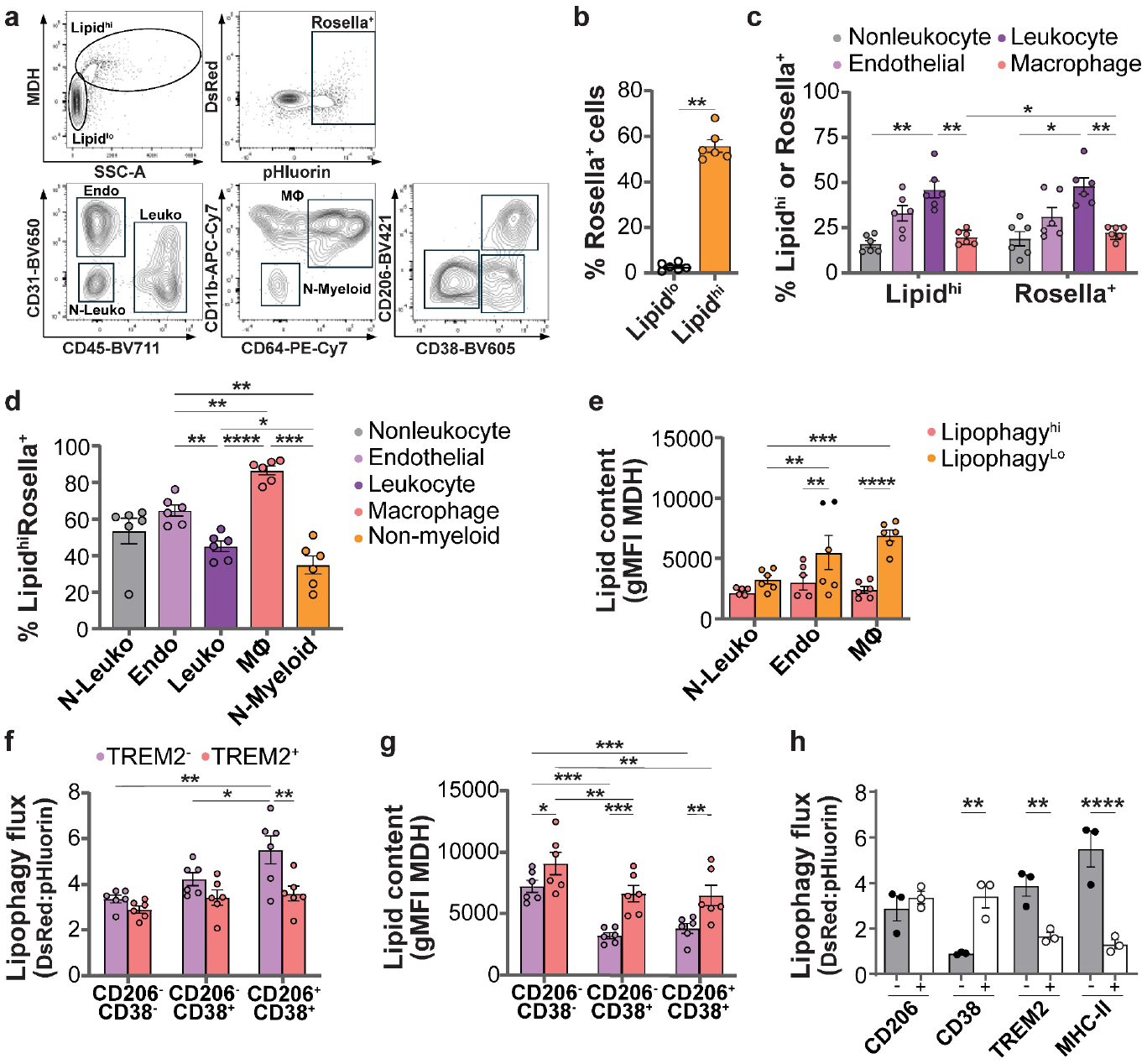
Lipophagy is impaired in TREM2-expressing macrophage foam cells of the atherosclerotic plaque. (**a**) Flow cytometry analysis of foamy (Lipid^hi^) and non-foamy (Lipid^lo^) cell populations, including Rosella^+^ cells, endothelial (Endo), leukocytes (Leuko), macrophages (MΦ) and non-myeloid leukocytes (N-myeloid) in aortic arches from hypercholesterolemic Rosella-PLIN2 mice. (**b**) Percentage of Rosella^+^ cells among Lipid^hi^ and Lipid^lo^ cells in aortic arches analysed in (**a**). (n=3 mice per sex pooled) (**c**) Percentage of indicated cell types among Lipid^hi^ or Rosella^+^ cells in aortic arches analysed in (**a**). (**d**) Percentage of Rosella^+^ cells among Lipid^hi^ cells in aortic arches analysed in (**a**). (**e**) Geometric mean fluorescence intensity (gMFI) of MDH in cells with high or low lipophagy flux among Rosella^+^ cell types identified in (**a**). (**f-g**) gMFI of DsRed:pHluorin ratio (**f**) and MDH (**g**) in Rosella^+^ plaque MΦ expressing (+) or not (-) indicated surface markers. (**h**) gMFI of DsRed:pHluorin ratio in thioglycolate-elicited peritoneal MΦ isolated from hypercholesterolemic Rosella-PLIN2 mice, expressing (+) or not (-) indicated surface markers. (n=3 female mice). Data are mean ± SEM, two-way ANOVA with Sidak post-hoc multiple comparison test.

Next, we assessed the lipid content of Rosella^+^ cells categorized based on high or low DsRed:pHluorin fluorescence ratios, designating them as Lipophagy^hi^ or Lipophagy^lo^, respectively. Lipophagy^lo^ macrophage and endothelial foam cells contained significantly more neutral lipids than their Lipophagy^hi^ counterparts (**Fig.6e**). No significant difference in lipid content was observed between Lipophagy^hi^ and Lipophagy^lo^ CD31^-^CD45^-^ foam cells, suggesting impaired lipophagy (**Fig.6e**).

To determine if macrophage subpopulations with distinct inflammatory and metabolic profiles exhibit variations in lipophagy flux, we compared lipophagy flux and lipid content in Rosella^+^ macrophage foam cells expressing or lacking surface markers of inflammation such as CD206, CD38, and TREM2, which have previously been linked to macrophage function in atherosclerotic lesions^22,32–35^. TREM2^+^ macrophage foam cells exhibited a higher lipid content than TREM2^-^ cells but had significantly lower lipophagy flux (**Fig.6f-g**). In contrast, CD206^+^ and CD38^+^ macrophages had higher lipophagy flux and lower lipid content, especially among TREM2^-^ macrophages. These findings indicate that macrophage lipophagy is highly heterogeneous and tightly regulated within atherosclerotic lesions.

To test if macrophage lipophagy heterogeneity is specific to the atheroma microenvironment, we elicited the recruitment of macrophage foam cells to the peritoneum of hypercholesterolemic Rosella-PLIN2 mice using thioglycolate, as previously described^36^. Consistent with our lesion data, lipophagy flux in TREM2^+^ and MHC-II^+^ macrophages was significantly lower compared to their TREM2^-^ and MHC-II^-^ counterparts, while CD38^+^ macrophages showed a much higher lipophagy flux (**Fig.6h**). These results suggest that TREM2^+^, CD38^-^ and MHC-II^+^ pro-inflammatory macrophage foam cells exhibit impaired lipophagy, accompanied by lipid accumulation, regardless of tissue environment. Overall, our findings highlight the dynamic lipid storage behavior of VSMC-derived foam cells in atherosclerosis and demonstrate the utility of the Rosella-PLIN2 model for studying LD formation and lipophagy flux *in vivo* across diverse foam cell populations.

## Discussion

Autophagy is essential for regulating macrophage functions such as efferocytosis, cholesterol efflux, and inflammasome activation – key processes for cardiovascular health. In atherosclerosis, marked by lipid accumulation and inflammation in the arterial wall, impaired autophagy is observed as a critical event that accelerates disease progression^6,16,37^. The crucial role of macrophage LD metabolism in atherosclerosis is emphasized by studies showing reduced foam cell lipid burden and plaque size in mice lacking whole-body or bone marrow-specific expression of the LD-associated protein PLIN2^38^. Similarly, organelle-specific autophagic pathways are gaining attention for their targeted impact on cardiometabolism^17,39,40^. However, the regulation of lipophagy, the selective autophagic degradation of LDs, remains poorly understood, especially *in vivo*.

In this study, we investigated the role of lipophagy in regulating foam cell immunometabolism. Using a novel Rosella-PLIN2 fluorescent lipophagy reporter, we successfully tracked lipophagy in human THP-1 macrophages and for the first time *in vivo* in a mouse model of atherosclerosis. We found that mTOR inhibition induced macrophage lipophagy, but that this effect could be blocked by disrupting the autophagic machinery or knocking down key lipophagy factors (VPS4A, SPG20, ORP8)^13–15^. Furthermore, stimulation of lipophagy using LD-ATTECs or αGFP-LC3 chimeric nanobodies enhanced LD degradation, mitochondrial respiration and cholesterol efflux in macrophage foam cells, supporting the idea that lipophagy plays a central role in macrophage lipid metabolism, independent of other autophagy pathways^11^.

Autophagy is modulated by cytokine- and TLR-mediated macrophage activation, influencing their inflammatory polarization^23,24,41^. Given that activated macrophages generate LDs^25,26,42^, we hypothesized that lipophagy contributes to macrophage polarization. Indeed, lipophagy flux was upregulated in TLR2-stimulated and M1 macrophages, but not in M2 macrophages. Interestingly, lipophagy stimulation mitigated CD80 upregulation in M1 macrophages, suggesting that lipophagy provides negative feedback on pro-inflammatory activation. Conversely, stimulating lipophagy during M2 polarization enhanced CD206 expression, consistent with the observation that lysosomal lipolysis sustains M2 activation^42^. Together, this data highlights the role of lipophagy in regulating macrophage foam cell immunometabolism, a novel therapeutic avenue for the treatment of atherosclerosis^43,44^.

Research on LD metabolism in arterial foam cells has been limited by the lack of lipophagy-specific tools and factors^17^. To track LDs and lipophagy flux *in vivo*, we used the Rosella-PLIN2 biosensor in C57BL/6N mice by tagging endogenous *Plin2* with Rosella, similarly to tdTom-Plin2 knock-in mice^29^. This allowed the fluorescent tagging of LDs and quantification of lipophagy flux, and we further investigated the regulation of LD metabolism and lipophagy flux in arterial foam cell populations during atherosclerosis by inducing hypercholesterolemia in Rosella-PLIN2 mice. Using CD68 and CD45 markers, we identified macrophage and VSMC foam cells and quantified PLIN2 expression. We found that macrophage foam cells were highly enriched in PLIN2, while only half of VSMC foam cells were PLIN2^+^. In line with previous work^8^, this supports the existence of a VSMC foam cell population that accumulates lipids outside of LDs, presumably inside LAL-defective lysosomes. The phenotypic differences between these newly identified subpopulations of VSMC foam cells may extend to other key cellular features, such as contractility or proliferative capacity.

Compared to macrophages, VSMCs have lower basal autophagy flux and impaired cholesterol efflux due to reduced ABCA1 expression^4^. However, pharmacological activation of autophagy with metformin increased ABCG1-mediated efflux to HDL^6^. Additionally, treatment of cholesterol-loaded VSMCs with HDL or apoA1 abolishes their macrophage-like phenotype, further supporting the atheroprotective potential of cholesterol efflux^45^. Here, stimulating lipophagy flux in primary VSMC foam cells with LD-ATTECs reduced their LD burden, suggesting that despite its low basal activity, lipophagy could be boosted to promote LD degradation in PLIN2^+^ arterial VSMC foam cells.

Consistent with our aortic sinuses data, flow cytometry analysis of aortic digests from atherosclerotic Rosella-PLIN2 mice revealed that almost all non-foam cells lacked Rosella, while about half of endothelial- and VSMC-like foam cells and 85% of macrophage foam cells expressed Rosella-PLIN2. Lipophagy was inversely correlated with lipid content in macrophage and endothelial foam cells, but not in VSMCs, suggesting a VSMC-specific defect in lipophagy. This finding aligns with our previous study reporting autophagy flux defects in VSMCs compared to macrophages during atherosclerosis progression^6^. LDs were recently implicated in endothelial dysfunction and inflammation, with neutral lipolysis in endothelial cells reported as being atheroprotective^46–48^. While confirming that arterial endothelial cells form *bona fide* LDs, our results point to endothelial lipophagy as a promising therapeutic target to prevent endothelial dysfunction and inflammation driving atherogenesis.

Emerging evidence identifies TREM2 as marker of a non-inflammatory, foamy macrophage subset in atherosclerotic plaques^49^, and recent findings suggest it regulates foam cell formation and survival^32,50^. In our Rosella-PLIN2 aortic digests, TREM2^+^ macrophages displayed high lipid content and low lipophagy, suggesting a lipophagy defect that may contribute to their lipid burden. This is consistent with observations that TREM2 signaling inhibits autophagy via mTOR^51^, potentially promoting foam cell formation in atherosclerosis. Our *in vitro* data indicate an interplay between lipophagy and macrophage polarization. Further analysis of arterial foam cell subsets based on CD38 (an M1-like marker) and CD206 (an M2-like marker) expression revealed that TREM2^-^ macrophages expressing CD38 and CD206 had low lipid content and high lipophagy. Notably, we identified a TREM2^-^CD206^-^CD38^-^ subpopulation with altered LD metabolism like TREM2^+^ macrophages. Interestingly, CD38^+^ macrophages, despite containing LDs, wouldn’t typically be classified as foam cells due to their lipid content. CD38, as a target of LXR, regulates cholesterol metabolism within macrophage foam cells. Its deficiency or inhibition leads to increased cholesterol accumulation in lysosomes, exacerbating atherosclerosis^52,53^. Notably, *Cd38* was one of the most significantly downregulated genes in macrophage foam cells from *Ldlr*^*-/-*^*Trem2*^*-/-*^ atherosclerotic mice, compared to *Ldlr*^*-/-*^*Trem2*^*+/+*^ mice^50^. This positions CD38 as potential marker of lipophagy-efficient macrophage foam cells and a regulator of cholesterol efflux during atherogenesis.

In conclusion, our findings reveal diverse foam cell subpopulations with heterogenous LD metabolism in atherosclerotic plaques, underscoring the need for targeted strategies in developing lipid metabolism-modulating therapies. The Rosella-PLIN2 mouse, with its ubiquitous PLIN2 expression, presents a promising tool for advancing research on LDs and lipophagy in cardiovascular diseases and beyond. This study highlights the critical role of lipophagy in macrophage immunometabolism, showing how macrophage foam cell LD degradation drives lipid metabolism and inflammatory profiles, offering new insights into atheroprotective mechanisms.

## Methods

### Generation of Rosella-Plin2 mice

All procedures on animals at The Centre for Phenogenomics (TCP) were reviewed and approved by TCP’s Animal Care Committee. TCP is certified by the Canadian Council on Animal Care and registered under the Animals for Research Act of Ontario. The *Plin2*^*em1Tcp*^ allele (MGI:7489806), here called Rosella-PLIN2, has the Rosella open reading frame inserted between bases 86586916 and 86586917 of Chr4 (GRCm39). The Rosella-PLIN2 mouse line was made at The Centre for Phenogenomics (Toronto, ON, Canada) using adeno-associated virus to deliver the repair template^54^ and electroporation to deliver Cas9 ribonucleoprotein^55^. The repair template consisted of 500-bp homology arms flanking the Rosella cDNA^19^ between AAV ITRs and was synthesized and packaged into AAV by VectorBuilder (Chicago, IL, USA). On the day of embryo manipulation, 1200 ng/µL of Cas9 protein (Integrated DNA Technologies, 1074182) was complexed with 378 ng/µL *Plin2* sgRNA (5’-AGCAGTAGTGGATCCGCAAC-3’) made using the EnGen sgRNA synthesis kit (New England Biolabs, E3322) in 1X Cas9 buffer (100 mM KCl, 20 mM Hepes, pH 7.2-7.4) for 10 minutes at 37ºC and kept on ice until electroporation. C57BL/NCrl (Charles River Laboratories, Strain code 027) mouse zygotes were collected from superovulated and mated females, treated with hyaluronidase to remove cumulus masses^56^, then treated briefly with acid Tyrode’s^55^, and placed in pre-equilibrated KSOM^AA^ (Zenith Biotech, ZEKS-50) containing ∼10^8^ AAV/µL. Zygotes were co-incubated with AAV for four hours at 37ºC with 6% CO_2_ prior to being washed and placed in a 1:1 mixture of Cas9 RNP and Opti-MEM (ThermoFisher 31985062). Zygotes were electroporated with 12 x 30 V 1-msec pulses with 100 msec intervals. Electroporated zygotes were returned to KSOM^AA^ media with AAV and incubated at 37ºC with 6% CO2 overnight. Two-cell embryos were washed and then transferred into CD-1 (Charles River Labs, Strain code 022) surrogate host mothers for gestation and birth.

Founders with the *Rosella* insertion were identified among born pups using genomic DNA for PCR with primers flanking the sgRNA target site and from inside the Rosella CDS to outside the 5’ homology arm, before quality control (QC). Template copy number was assessed by real-time PCR (ThermoFisher, Viia7) using a known single-copy DsRed knock-in mouse as a calibrator. Founders without extra template copies were then assessed by long-range PCR followed by long-read sequencing of PCR amplicons (Plasmidsaurus, San Francisco, CA, USA) to confirm site-specific integration and sequence integrity. Four founders that passed QC were bred to C57BL/6NCrl mice. Born pups were identified with the same PCRs and subjected to the same quality control as founders. N1 mice that passed QC were used to establish the Rosella-PLIN2 mouse line by intercrossing.

### Animal experiments

All animal procedures were approved by the University of Ottawa Animal Care and Use Committee. Male and female C57BL/6NCrl or Rosella-PLIN2 mice were used for primary cell culture experiments and to monitor LD dynamics in foam cells during atherosclerosis development. Male and female mice aged 8 to 10 weeks were injected with 5×10^11^ particles of AAV-PCSK9 (University of Pennsylvania) in the peritoneum as previously and fed a Western Diet (Envigo, TD.88137) for 12, 16 or 22 weeks. The mice were randomly assigned to each of the cohorts as described. For the analysis of foamy peritoneal macrophages, 12-week-fed Rosella-PLIN2 mice were injected intraperitoneally with 1mL of 3% thioglycolate (BD Difco) 3 days before endpoint. The mice were anesthetized by isoflurane inhalation and euthanized by exsanguination. Blood was collected by cardiac puncture and mice were perfused with HBSS.

### Plasmids

pLenti-Ef1a-Rosella-hPLIN2 was generated at the uOttawa Genomic Editing and Molecular biology (GEMb) facility. The coding sequence of hPLIN2 was amplified from a vector in the MGC Human ORFeome collection (Transomic Technologies) using oligonucleotides with sequence 5’–AAATCTGATCACATGGCATCCGTTGCAGTTGA-3’ and 5’– TGATTATCATATGACTAGTCCCGGGCTAATGAGTTTTATGCTCAGATCGCTGG-3’. The Rosella dual fluorescent protein fusion construct was amplified from pAS1NB c Rosella I (Addgene #71245)^19^ using primers 5’-ACTAGCCTCGAGGTTTAAACTACGGGCCACCATGGCCTCCTC-3’ and 5’– CAACGGATGCCATGTGATCAGATTTGTATAGTTCATCCATGCCT-3’). These two PCR amplicons were then inserted into EcoRI and BamHI cut pWPXLd by Gibson assembly^57^ such that Rosella-hPLIN2 fusion expression is driven by the EF1a promoter. Inducible constructs expressing the GFP-nanobody alone or fused to LC3B (pInducer-GFPnano-Ctrl/LC3B)^21^ were generated by PCR amplification of GFP-nanobody only and GFP-nanobody fusion constructs in pRGZCV using the following primer pair: 5’-CGCGGCCCCGAACTAGTCCAGTGTGGCGGCCGCGGATCCGCCG-3’ and 5’– CTAGACTCGAGCGGCCGCCACTGTGGGGGCGGAATTTGCGGCCGC-3’, which were combined in equimolar ratios with BstXI cut pINDUCER20^58^ in an isothermal DNA assembly reaction. Lentivirus particles were generated by co-transfection of pLenti-Ef1a-Rosella-hPLIN2 or plasmids pInducer-GFPnano-Ctrl/LC3B with psPAX2 and pMD2.G (Addgene) in HEK293FT cells (ThermoFisher Scientific) using Lipofectamine 3000 transfection reagent. Supernatants were collected 48 hours post-transfection, and lentiviral particles were concentrated using Lenti-X Concentrator (Takara Bio).

### Cell culture

Bone marrow was isolated by perfusion of femurs and tibias from 7- to 12-week-old C57BL/6N mice and cultured in DMEM (Gibco) with 10% heat-inactivated fetal bovine serum (FBS, Gibco), 2 mM GlutaMAX, 20% L929-conditioned medium, 100 U/mL penicillin, and 100 µg/mL streptomycin (Pen-Strep) for 7 days, with media change on day 3. After differentiation, BMDMs were replated in DMEM with 10% FBS, 10% L929-conditioned medium and Pen-Strep (BMDM media). Peritoneal macrophages were harvested from 8-week-old male and female C57BL/6N mice 3 days after intraperitoneal injection of 1 ml 3% thioglycolate (BD Difco) and cultured in DMEM with 10% FBS and 1% Pen-Strep. Primary smooth muscle cells were isolated from the thoracic aorta of C57BL/6N or Rosella-PLIN2 mice. The aorta was digested for 20-30 minutes at 37°C (2U/mL of Liberase TM, 2U/mL of Elastase [Worthington Biochemical] in HBSS), as previously described^6^.Cells were plated on 0.1% gelatin coated plates for the first passage, and cultured in DMEM supplemented with 20% FBS, 1% Antibiotic-Antimycotic (Gibco), 1x L-glutamine, 10ng/mL of recombinant mouse leukemia inhibitory factor (LIF; PeproTech) and 0.1mM 2-mercaptoethanol (Gibco).

THP-1 and HEK293T cells were purchased from and authenticated by ATCC and confirmed mycoplasma-negative before experiments. THP-1 and HEK293T were respectively cultured in RPMI-1640 (Gibco) and DMEM supplemented with 10% FBS and Pen-Strep. THP-1 monocytes were differentiated into macrophages by incubation with 25 ng/mL PMA for 72 hours. Where specified, macrophages were transfected in OptiMEM media Human On-Target Plus siRNA SmartPools (Dharmacon) using Lipofectamine RNAiMAX (Invitrogen) according to the manufacturer’s instructions. After 24h, an equal volume of THP-1 media was added, and cells were incubated for another 24h before lipid loading and treatments. Stable THP-1 cell lines (Rosella-PLIN2 and doxycycline-inducible αGFP-Ctrl and αGFP-LC3B) were generated by lentiviral transduction of monocytes in the presence of 8 µg/ml of polybrene. Rosella-PLIN2 cells were selected by fluorescence-activated cell sorting using the BD FACS Aria IIIu (Beckton Dickinson), while αGFP-Ctrl and αGFP-LC3B THP-1 cell lines were selected with 500 µg/ml of G418.

### Lipid loading and treatments

All macrophages were lipid-loaded by incubation with 50 µg/ml of agLDL for 24 hours. Where indicated, differentiated THP-1 macrophages and HEK293T cells were lipid-loaded with 200µM of oleic acid (OA) pre-complexed with fatty acid-free BSA (Sigma) at a 1:3 ratio. VSMCs were lipid-loaded with 35μg/mL of cholesterol from mβ-CD-cholesterol complexes (Sigma) for 48 hours in DMEM media supplemented with 10% FBS and 1% P/S. Where indicated, lipid-loaded cells were washed and incubated for 24h in DMEM supplemented with 2 mg/ml of fatty acid-free BSA (Sigma). During this incubation, cells were treated with Rapamycin (100 nM, Sigma), Torin-1 (250 nM, Cayman), LD-ATTEC4 (15 µM, synthesized as previously described^20^ by Novalix, France) or Doxycycline (100 ng/mL), and, where specified, in presence of 50 μg/mL of human recombinant ApoA1 or HDL (generated as previously^59^).

For autophagy flux analyses, cells were treated for 2 h prior imaging or cell lysis with either chloroquine (30 µM, Sigma), Bafilomycin A1 (75 nM, Invivogen) or vehicle control (DMSO). For TLR stimulations, THP-1 macrophages were treated for 24h in THP-1 media with LPS (100 ng/mL, O111:B4 Sigma), Pam3Csk4 (500 ng/mL, Invivogen) or Lipoteichoic acid (100 ng/mL, LTS, Invivogen). For macrophage polarizations, cells were incubated for 24h with 100 ng/mL of LPS plus 20 ng/mL of murine (PreproTech) or human (R&D Systems) IFNγ (M1), or with 20 ng/mL of murine IL-4 (PreproTech, M2 BMDMs) or 20 ng/mL of human IL-4 plus IL-13 (PreproTech, M2 THP-1).

### Cell metabolism

Macrophages cultured and treated in Seahorse 96-well plates were washed twice and incubated in XF DMEM or RPMI assay medium (Agilent) supplemented with 11 mM XF glucose (Agilent), 1mM XF pyruvate (Agilent) and 2mM XF glutamine (Agilent) for 1 hour at 37°C without CO_2_. Oxygen Consumption Rate (OCR) and Extracellular Acidification Rate (ECAR) were measured using the Seahorse XF Mito Stress Test Kit on the XFe96 analyzer according to the manufacturer’s instructions. Inhibitors were added at the following final concentrations: 1 µM of oligomycin, 3 µM of FCCP, 0.5 µM of Rotenone and 1 µM of antimycin A (all from Sigma). At the end of the measurements, live Hoechst 33342 (Invitrogen) was added to the cells to stain the nuclei, and cells were imaged using the Cytation 5 automated widefield microscope. Cell nuclei were counted using the BioGen 5 software to normalize OCR and ECAR measurements.

### Cholesterol efflux

Macrophages were loaded as previously described^59^ with 50μg/mL agLDL and 0.5μCi/mL ^3^H-cholesterol (NET139001MC, Perkin-Elmer) in BMDM or THP-1 media for 24 hours, followed by two washes in DPBS and an overnight incubation in DMEM with 2mg/mL fatty acid-free BSA. Cholesterol efflux to 50μg/mL human recombinant apoA1 or HDL in the same media was carried out after 24 hours in the presence of 15μM of LD-ATTECs, 100 ng/mL of doxycycline, or vehicle control (DMSO). Supernatants were then removed, and cells were lysed in 0.5M NaOH. Cell and supernatant radioactivity were quantified using the Hydex Sense plate reader (Gamble). Cholesterol efflux was calculated as a percentage of ^3^H-cholesterol in the supernatant/(^3^H-cholesterol in the supernatant + ^3^H-cholesterol in cells)x100%.

### Immunostaining

VSMCs and BMDMs cultured on ibiTreat 18 well chamber slides (Ibidi) were accordingly treated, fixed with 4% paraformaldehyde in PBS (ThermoFisher Scientific) at 37°C for 10 minutes, and washed five times with 1X PBS (ThermoFisher Scientific). After blocking and permeabilizing in 5% BSA with 0.1% Triton-X for 30 minutes at room temperature (RT), cells were incubated overnight at 4°C with anti-PLIN2 (ab52356; 1:400) in 1% BSA with 0.05% saponin. Cells were then incubated with an anti-Rabbit IgG Alexa Fluor Plus 647 secondary antibody (1:1000) in 1% BSA with 0.05% saponin for 2h at RT. Cells were washed three times in 1X PBS between all antibody incubations. Cells were counterstained with BODIPY 493/503 (Invitrogen) for 30 minutes at RT and with DAPI (Invitrogen) for 5 minutes at RT, then washed and left in 1X PBS for imaging.

Mouse hearts were embedded in optimal cutting temperature compound (OCT), flash frozen in liquid nitrogen, and stored at -80C until cryosectioning. The embedded aortic sinuses were serial sectioned at a 10µm thickness, and stored at - 20°C. Sinuses were fixed for 10 minutes at RT with 4% PFA (or 3.7% PFA in HEPES pH 7 as described previously^60^ for Rosella-PLIN2 hearts) and blocked/permeabilized with 10% Normal Horse Serum (NHS) with 0.1% Triton-X 100 in PBS 1X for 30 minutes at RT. Then primary antibodies (Table 1) were diluted in 1% NHS in PBS 1X and incubated overnight on sinuses at 4°C in a moisture chamber. Secondary antibodies (Table 1) were diluted in 1% NHS in PBS 1X and incubated for 1h at RT in a moisture chamber. The primary conjugated anti-CD68 antibody (Novus, NPB2-33337AF750 at 1:100) was diluted in 1% NHS and incubated for 2h at RT in a moisture chamber. All antibody steps were followed by three washes in 1X PBS. The sections were counterstained with DAPI for 5 minutes at RT, washed and mounted on #1.5 coverslips with Prolong Glass Antifade Mountant (Invitrogen). Sinuses were imaged using the inverted Zeiss AxioObserver 7 with the Zen Pro 3.1 software and using the 20x objective. Image processing and image analysis were done using the Zen Pro 3.1 software.

### Flow cytometry

THP-1 macrophages were detached with enzyme-free Cell Dissociation Buffer (Gibco) at 4°C and resuspended in FACS buffer (DPBS with 2% FBS and 1mM EDTA). To analyze atheroma foam cells, thoracic arches from atherosclerotic mice were dissected, cleaned and digested with 0.4 units/mL Liberase TM (Roche), 40 units/mL hyaluronidase (Sigma), and 20 units/mL DNAse (Sigma) in HBSS for 15-20 minutes at 37°C on a rotator. The resulting cell suspension was filtered through a 70μm cell strainer and resuspended in FACS buffer. Peritoneal cells were isolated by lavage of thioglycolate-injected mice with cold DPBS, followed by red blood cell lysis with PharmLyse buffer 1X (BD Biosciences) and resuspension in FACS buffer. For antibody staining, cells were incubated with human or murine FcBlock (BD Biosciences), then with conjugated antibodies (Table 1) for 30 min at 4°C. Afterward, cells were stained with Fixable Viability Stain 700 (1:1000) and Monodansylpentane (MDH, Abcepta; 1:1000) for 30 minutes at 4°C, washed in FACS buffer and acquired on a 5-laser Cytek Aurora spectral flow cytometer (Cytek Biosciences). Data were analyzed using SpectroFlo 3.3.0 (Cytek Biosciences) and FlowJo 10.8.2 (BD Biosciences).

### qPCR

Cells were lysed in Trizol (Ambion), and RNA was isolated using the Direct-zol RNA MiniPrep kit (Zymo). Reverse transcription was carried out with iScript Reverse Transcription Supermix (Bio-Rad). qPCR was performed using the CFX Connect Real Time PCR detection System (Bio-Rad). Primer sequences are listed in Table 2. mRNA levels were quantified using the comparative CT method, with *HPRT1* as the housekeeping gene.

### Western Blotting

Cells were lysed in 2X Laemmli Sample Buffer (Bio-Rad) containing β-mercaptoethanol and boiled at 95°C for 5 min. Samples were run on 8%–16% Criterion TGX Stain Free Pre-cast Gels (BioRad) and UV-activated for total protein quantification using the ChemiDoc XRS + System (Bio-Rad). Proteins were transferred onto 0.22μm PVDF Membranes (Bio-Rad) using the Trans-Blot Turbo Transfer System (Bio-Rad). Immunoblotting for indicated proteins was performed with primary antibodies (Table 1) overnight at 4°C followed by horseradish peroxidase conjugated secondary antibodies for 1 h at room temperature. Proteins were developed using either Clarity (Bio-Rad) or Clarity Max (Bio-Rad) ECL Substrates and imaged on the ChemiDoc XRS + system (Bio-Rad).

### Cell Imaging

For live-cell microscopy, cells were cultured on ibiTreat 18-well chamber slides (Ibidi) and stained for 30 minutes at 37°C with MDH (1:10,000) or LysoTracker Deep Red (50nM, Invitrogen) in FluoroBrite DMEM (Gibco) (BMDMs and VSMCs) or phenol red-free RPMI (Gibco) (THP-1) with 2 mg/mL fatty acid-free BSA before imaging. Live or fixed cells were imaged on the Zeiss LSM880 confocal microscope with Airyscan, using the 20x, 0.8 NA, Air, Plan-Apochromat or 63x, 1.4 NA, Oil, Plan-Apo (NA objectives) or on the Zeiss AxioObserver 7 inverted widefield microscope with the 20x 0.8 NA Plan-Apochromat or 63x 1.4 NA, Oil, Plan-Apochromat objectives, at 37°C and 5% CO_2_ or RT. To assess lipophagy flux at the single-cell level, we quantified lipophagy events per cell using the mito-QC Counter^61^, or the DsRed:pHluorin fluorescence ratio per cell. Lipophagy events, LD area, and DsRed:pHluorin fluorescence ratios per cell were analyzed in Image J. For cells imaged live on the widefield Cytation imaging reader (BioTek), with a 20x objective at 37°C and 5% CO_2_, images were analyzed using the BioGen5 software.

### Statistical analyses

Data are presented as mean ± SEM or mean ± 95%CI and represent at least 3 independent experiments, unless otherwise noted. Statistical analysis was performed using Prism v10.2.0 (GraphPad Software Inc) and are detailed in the figure legends. Group differences were assessed using two-tailed unpaired Student’s t test, one-way or two-way ANOVA with Tukey’s, Dunnett’s, Sidak’s or Holm-Sidak’s multiple comparison corrections. Statistical significance is indicated in the figures as *p < 0.05, **p < 0.01, ***p < 0.001, ****p < 0.0001.

## Data availability

The data supporting the findings of this study are available in the article file and its supplementary information.

## Acknowledgements

pAS1NB c Rosella I was a gift from Mark Prescott (Addgene plasmid # 71245 ; http://n2t.net/addgene:71245 ; RRID:Addgene_71245). pWPXLd was a gift from Didier Trono (Addgene plasmid # 12258; http://n2t.net/addgene:12258; RRID:Addgene_12258). pInducer20 was a gift from Stephen Elledge (Addgene plasmid # 44012; http://n2t.net/addgene:44012 ; RRID:Addgene_44012). All the lentivectors were generated at the Genomic Engineering and Molecular Biology (GEMb) core facility in the Faculty of Medicine at the University of Ottawa (RRID:SCR_022954). We thank the University of Ottawa Animal Care and Veterinary Services for their assistance with the animal experiments. We thank Chloe van Oostende-Triplet from the uOttawa Cell Biology and Image Acquisition Core Facility and Vera A. Tang from the uOttawa Flow Cytometry and Virometry Core Facility for their help with the microscopy and flow cytometry analysis, respectively. We thank Gregory Fairn (St Michael’s Hospital) for providing us with the recombinant human apoA1 plasmid. Illustrations were made using BioRender. This work was supported by the Canadian Institutes for Health Research (PJT-391187 and Canada Research Chair to M.O., the Heart and Stroke Foundation of Canada (M.O.), the Canada Graduate Scholarship-Master’s (V.R.), the Vanier Canada Graduate Scholarship (D.B.) and the UOHI Endowed Fellowship (T.L., V.R. and V.L.).

## Author contributions

T.L. and M.O. conceived the study and designed the experiments; T.L., N.J., V.R., D.B., C.E., V.L., M.N. and M.G. performed experiments; T.L., N.J., V.R. and D.B. analyzed the data; L.M.J.N. generated and provided the Rosella-PLIN2 reporter mouse line; T.L. and M.O. wrote the original manuscript, with input from all authors; M.O. secured funds; K.J.R., D.G. and R.C.R. provided resources; T.L., R.C.R. and M.O. supervised the project.

## References

1. Libby, P. The changing landscape of atherosclerosis. Nature 592, 524–533 (2021).

2. Moore, K. J., Sheedy, F. J. & Fisher, E. A. Macrophages in atherosclerosis: a dynamic balance. Nat Rev Immunol 13, 709–721 (2013).

3. Allahverdian, S., Chehroudi, A. C., McManus, B. M., Abraham, T. & Francis, G. A. Contribution of intimal smooth muscle cells to cholesterol accumulation and macrophage-like cells in human atherosclerosis. Circulation 129, 1551–1559 (2014).

4. Wang, Y. et al. Smooth Muscle Cells Contribute the Majority of Foam Cells in ApoE (Apolipoprotein E)-Deficient Mouse Atherosclerosis. Arterioscler Thromb Vasc Biol 39, 876–887 (2019).

5. Feil, S. et al. Transdifferentiation of vascular smooth muscle cells to macrophage-like cells during atherogenesis. Circ Res 115, 662–667 (2014).

6. Robichaud, S. et al. Autophagy Is Differentially Regulated in Leukocyte and Nonleukocyte Foam Cells During Atherosclerosis. Circ Res 130, 831–847 (2022).

7. Shankman, L. S. et al. KLF4-dependent phenotypic modulation of smooth muscle cells has a key role in atherosclerotic plaque pathogenesis. Nat Med 21, 628–637 (2015).

8. Dubland, J. A. et al. Low LAL (Lysosomal Acid Lipase) Expression by Smooth Muscle Cells Relative to Macrophages as a Mechanism for Arterial Foam Cell Formation. Arterioscler Thromb Vasc Biol 41, e354–e368 (2021).

9. Ouimet, M., Barrett, T. J. & Fisher, E. A. HDL and Reverse Cholesterol Transport. Circ Res 124, 1505–1518 (2019).

10. Ouimet, M. et al. Autophagy regulates cholesterol efflux from macrophage foam cells via lysosomal acid lipase. Cell Metab. 13, 655–667 (2011).

11. Robichaud, S. et al. Identification of novel lipid droplet factors that regulate lipophagy and cholesterol efflux in macrophage foam cells. Autophagy 17, 3671– 3689 (2021).

12. Singh, R. et al. Autophagy regulates lipid metabolism. Nature 458, 1131–1135 (2009).

13. Pu, M. et al. ORP8 acts as a lipophagy receptor to mediate lipid droplet turnover. Protein & Cell pwac063 (2022) doi:10.1093/procel/pwac063.

14. Das, D. et al. VPS4A is the selective receptor for lipophagy in mice and humans. Mol Cell 84, 4436-4453.e8 (2024).

15. Chung, J. et al. The Troyer syndrome protein spartin mediates selective autophagy of lipid droplets. Nat Cell Biol 25, 1101–1110 (2023).

16. Razani, B. et al. Autophagy links inflammasomes to atherosclerotic progression. Cell Metab 15, 534–544 (2012).

17. Laval, T. & Ouimet, M. A role for lipophagy in atherosclerosis. Nat Rev Cardiol 20, 431–432 (2023).

18. Sztalryd, C. & Brasaemle, D. L. The perilipin family of lipid droplet proteins: Gatekeepers of intracellular lipolysis. Biochim Biophys Acta Mol Cell Biol Lipids 1862, 1221–1232 (2017).

19. Rosado, C. J., Mijaljica, D., Hatzinisiriou, I., Prescott, M. & Devenish, R. J. Rosella: a fluorescent pH-biosensor for reporting vacuolar turnover of cytosol and organelles in yeast. Autophagy 4, 205–213 (2008).

20. Fu, Y. et al. Degradation of lipid droplets by chimeric autophagy-tethering compounds. Cell Res 31, 965–979 (2021).

21. Rothbauer, U. et al. A versatile nanotrap for biochemical and functional studies with fluorescent fusion proteins. Mol Cell Proteomics 7, 282–289 (2008).

22. Dib, L. et al. Lipid-associated macrophages transition to an inflammatory state in human atherosclerosis increasing the risk of cerebrovascular complications. Nat Cardiovasc Res 2, 656–672 (2023).

23. Fang, L., Wu, H.-M.Ding, P.-S. & Liu, R.-Y. TLR2 mediates phagocytosis and autophagy through JNK signaling pathway in Staphylococcus aureus-stimulated RAW264.7 cells. Cell Signal 26, 806–814 (2014).

24. Xu, Y. et al. Toll-like receptor 4 is a sensor for autophagy associated with innate immunity. Immunity 27, 135–144 (2007).

25. Feingold, K. R. et al. Mechanisms of triglyceride accumulation in activated macrophages. J. Leukoc. Biol. 92, 829–839 (2012).

26. Huang, Y. et al. Toll-like receptor agonists promote prolonged triglyceride storage in macrophages. J. Biol. Chem. 289, 3001–3012 (2014).

27. Boucher, D. M., Vijithakumar, V. & Ouimet, M. Lipid Droplets as Regulators of Metabolism and Immunity. Immunometabolism 3, e210021 (2021).

28. Bosch, M. & Pol, A. Eukaryotic lipid droplets: metabolic hubs, and immune first responders. Trends Endocrinol Metab 33, 218–229 (2022).

29. Madsen, S. et al. A fluorescent perilipin 2 knock-in mouse model reveals a high abundance of lipid droplets in the developing and adult brain. Nat Commun 15, 5489 (2024).

30. Francis, G. A. The Greatly Under-Represented Role of Smooth Muscle Cells in Atherosclerosis. Curr Atheroscler Rep 25, 741–749 (2023).

31. Najt, C. P., Devarajan, M. & Mashek, D. G. Perilipins at a glance. J Cell Sci 135, jcs259501 (2022).

32. Patterson, M. T. et al. Trem2 promotes foamy macrophage lipid uptake and survival in atherosclerosis. Nat Cardiovasc Res 2, 1015–1031 (2023).

33. Fernandez, D. M. et al. Single-cell immune landscape of human atherosclerotic plaques. Nat Med 25, 1576–1588 (2019).

34. Cole, J. E. et al. Immune cell census in murine atherosclerosis: cytometry by time of flight illuminates vascular myeloid cell diversity. Cardiovasc Res 114, 1360– 1371 (2018).

35. Robichaud, S., Rochon, V., Emerton, C., Laval, T. & Ouimet, M. Trehalose promotes atherosclerosis regression in female mice. Front Cardiovasc Med 11, 1298014 (2024).

36. Li, A. C. et al. Differential inhibition of macrophage foam-cell formation and atherosclerosis in mice by PPARα, β/δ, and γ. J Clin Invest 114, 1564–1576 (2004).

37. Liao, X. et al. Macrophage autophagy plays a protective role in advanced atherosclerosis. Cell Metab 15, 545–553 (2012).

38. Paul, A., Chang, B. H.-J., Li, L., Yechoor, V. K. & Chan, L. Deficiency of adipose differentiation-related protein impairs foam cell formation and protects against atherosclerosis. Circ Res 102, 1492–1501 (2008).

39. Jeong, S.-J., Zhang, X., Rodriguez-Velez, A., Evans, T. D. & Razani, B. p62/SQSTM1 and Selective Autophagy in Cardiometabolic Diseases. Antioxid Redox Signal 31, 458–471 (2019).

40. Bravo-San Pedro, J. M., Kroemer, G. & Galluzzi, L. Autophagy and Mitophagy in Cardiovascular Disease. Circ Res 120, 1812–1824 (2017).

41. Liu, K. et al. Impaired macrophage autophagy increases the immune response in obese mice by promoting proinflammatory macrophage polarization. Autophagy 11, 271–284 (2015).

42. Huang, S. C.-C. et al. Cell-intrinsic lysosomal lipolysis is essential for alternative activation of macrophages. Nat. Immunol. 15, 846–855 (2014).

43. Koelwyn, G. J., Corr, E. M., Erbay, E. & Moore, K. J. Regulation of macrophage immunometabolism in atherosclerosis. Nat Immunol 19, 526–537 (2018).

44. Certo, M., Rahimzadeh, M. & Mauro, C. Immunometabolism in atherosclerosis: a new understanding of an old disease. Trends Biochem Sci 49, 791–803 (2024).

45. Vengrenyuk, Y. et al. Cholesterol loading reprograms the microRNA-143/145-myocardin axis to convert aortic smooth muscle cells to a dysfunctional macrophage-like phenotype. Arterioscler Thromb Vasc Biol 35, 535–546 (2015).

46. Boutagy, N. E. et al. Dynamic metabolism of endothelial triglycerides protects against atherosclerosis in mice. J Clin Invest 134, e170453 (2024).

47. Kim, B. et al. Endothelial lipid droplets suppress eNOS to link high fat consumption to blood pressure elevation. J Clin Invest 133, e173160 (2023).

48. Tan, Y. et al. Lipid droplets sequester palmitic acid to disrupt endothelial ciliation and exacerbate atherosclerosis in male mice. Nat Commun 15, 8273 (2024).

49. Zernecke, A. et al. Integrated single-cell analysis-based classification of vascular mononuclear phagocytes in mouse and human atherosclerosis. Cardiovasc Res 119, 1676–1689 (2023).

50. Piollet, M. et al. TREM2 protects from atherosclerosis by limiting necrotic core formation. Nat Cardiovasc Res 3, 269–282 (2024).

51. Ulland, T. K. et al. TREM2 Maintains Microglial Metabolic Fitness in Alzheimer’s Disease. Cell 170, 649-663.e13 (2017).

52. Terao, R. et al. LXR/CD38 activation drives cholesterol-induced macrophage senescence and neurodegeneration via NAD+ depletion. Cell Rep 43, 114102 (2024).

53. Xu, X. et al. Lysosomal cholesterol accumulation in macrophages leading to coronary atherosclerosis in CD38(-/-) mice. J Cell Mol Med 20, 1001–1013 (2016).

54. Chen, S. et al. CRISPR-READI: Efficient Generation of Knockin Mice by CRISPR RNP Electroporation and AAV Donor Infection. Cell Rep 27, 3780-3789.e4 (2019).

55. Gertsenstein, M. & Nutter, L. M. J. Production of knockout mouse lines with Cas9. Methods 191, 32–43 (2021).

56. Behringer, R., Gertsenstein, M., Nagy, K. V. & Nagy, A. Manipulating the Mouse Embryo: A Laboratory Manual. (Cold Spring Harbor Laboratory Press, 2014).

57. Gibson, D. G. et al. Enzymatic assembly of DNA molecules up to several hundred kilobases. Nat Methods 6, 343–345 (2009).

58. Meerbrey, K. L. et al. The pINDUCER lentiviral toolkit for inducible RNA interference in vitro and in vivo. Proc Natl Acad Sci U S A 108, 3665–3670 (2011).

59. Robichaud, S. & Ouimet, M. Quantifying Cellular Cholesterol Efflux. Methods Mol Biol 1951, 111–133 (2019).

60. McWilliams, T. G. & Ganley, I. G. Investigating Mitophagy and Mitochondrial Morphology In Vivo Using mito-QC: A Comprehensive Guide. Methods Mol Biol 1880, 621–642 (2019).

61. Montava-Garriga, L., Singh, F., Ball, G. & Ganley, I. G. Semi-automated quantitation of mitophagy in cells and tissues. Mech Ageing Dev 185, 111196 (2020).

